# Protein structure determination using Riemannian approach

**DOI:** 10.1101/599761

**Authors:** Xian Wei, Zhicheng Li, Shijian Li, Xubiao Peng, Qing Zhao

**Affiliations:** Center for Quantum Technology Research, School of Physics, Beijing Institute of Technology, Beijing, China; Department of science, Taiyuan Institute of Technology, Taiyuan, China

## Abstract

The protein nuclear magnetic resonance (NMR) structure determination is one of the most extensively studied problems due to its increasing importance in biological function analysis. We adopt a novel method, based on one of the matrix completion (MC) techniques–the Riemannian approach, to solve the protein structure determination problem. We formulate the protein structure in terms of low-rank matrix which can be solved by an optimization problem in the Riemannian spectrahedron manifold whose objective function has been delimited with the derived boundary condition. Two efficient algorithms in Riemannian approach-the trust-region (Tr) algorithm and the conjugate gradient (Cg) algorithm are used to reconstruct protein structures. We first use the two algorithms in a toy model and show that the Tr algorithm is more robust. Afterwards, we rebuild the protein structure from the NOE distance information deposited in NMR Restraints Grid (http://restraintsgrid.bmrb.wisc.edu/NRG/MRGridServlet). A dataset with both X-ray crystallographic structure and NMR structure deposited in Protein Data Bank (PDB) is used to statistically evaluate the performance of our method. By comparing both our rebuilt structures and NMR counterparts with the “standard” X-ray structures, we conclude that our rebuilt structures have similar (sometimes even smaller) RMSDs relative to “standard” X-ray structures in contrast with the reference NMR structures. Besides, we also validate our method by comparing the Z-scores between our rebuilt structures with reference structures using Protein Structure Validation Software suit. All the validation scores indicate that the Riemannian approach in MC techniques is valid in reconstructing the protein structures from NOE distance information. The software based on Riemannian approach is freely available at https://github.com/xubiaopeng/Protein_Recon_MCRiemman.

**Author summary:** Matrix Completion is a technique widely used in many aspects, such as the global positioning in sensor networks, collaborative filtering in recommendation system for many companies and face recognition, etc. In biology, distance geometry used to be a popular method for reconstructing protein structures related to NMR experiment. However, due to the low quality of the reconstructed results, those methods were replaced by other dynamic methods such as ARIA, CYANA and UNIO. Recently, a new MC technique named Riemannian approach is introduced and proved mathematically, which promotes us to apply it in protein structure determination from NMR measurements. In this paper, by combining the Riemannian approach and some post-processing procedures together, we reconstruct the protein structures from the incomplete distance information measured by NMR. By evaluating our results and comparing with the corresponding PDB NMR deposits, we show that the current Riemannian approach method is valid and at least comparable with (if not better than) the state-of-art methods in NMR structure determination.

## Introduction

Three-dimensional protein structure plays a vital role in molecular conformation because of both the importance of the protein function and the applications on drug design and disease detection. Protein structures can be determined mainly through delicate experimental methods, including X-ray crystallography, nuclear magnetic resonance (NMR), Cryo-electron microscopy(cryo-EM) and so on. Different from single-crystal X-ray diffraction which has been largely automated, NMR spectroscopy requires skilled manual intervention. However, NMR spectroscopy is an important approach for measuring the 3D structure of proteins in solution under near physiological conditions [1].

The NMR method for protein structure measurement began in 1980s [2], and its spectroscopy provided a network of distance measurements between spatially proximate hydrogen atoms [3] [4]. The typical NMR-based protein structure determination pipeline involves peak picking from NMR spectra, chemical shift assignment (spectral assignment), assignment of geometric restraints and the structural calculation [5]. More specifically, this method has promoted a need for efficient computational algorithms.

One approach to obtain molecular conformations is related to the molecular dynamics and simulated annealing [6], such as ARIA [45], CYANA [7] and UNIO [8]. Another approach is to use distance geometry methods [9] where many algorithms such as EMBED [10], DISGEO [11], and DGSOL [12] [13] are proposed to interpret the macromolecular conformation based on NMR experimental data.

Recently another technology named matrix completion (MC) [14] [15] is a burgeoning topic drawing the attention of many researchers in the field of model reduction [16], pattern recognition [17] and machine learning [18]. This technology, an offshoot of compressed sensing (CS) [19] [20], seeks explicitly the lowest rank matrix consistent with the known entries by effective algorithms according to the dependence among matrix elements imposed by the low rank structure [21]. In particular, Jawanpuria et al. [22] whose group leads to a generic framework to the structured low-rank matrix learning problem have succeeded in solving the problem of learning a low-rank matrix through Riemannian approach. In the work of Jawanpuria et al. [22], the NP-hard rank minimization problem of MC is transferred into an optimization problem based on the Riemannian spectrahedron manifold. In their work, two efficient algorithms– conjugate gradient (Cg) and trust-region (Tr) are proposed and outperform other algorithms in robust.

In this paper, we treat the protein structure determination as a low-rank matrix completion problem since the protein structure can be formulated as a low-rank distance matrix. As a result, we can apply the algorithms of the Riemannian theory in determining the NMR-based protein structure. By taking those algorithms, we can avoid the high-dimensional problems and also provide a correct completed distance matrix for the determination.

## Methods

### Solution to determinate protein structure

The distance between all pairs of atoms in a large molecule can be transformed into a protein distance matrix *D* with nonnegative entries and zero diagonal [23], such that:

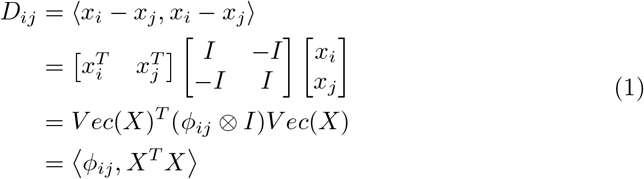

Where the corresponding coordinates are defined as *X*:= *x*_1_, *x*_2_,…, *x*_*n*_ ∈ ℜ in the three-dimensional cartesian coordinate system.

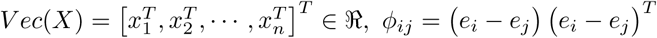, ⊗ denotes the Kronecker product.

Consider the Gram matrix *G*:= *X*^*T*^ *X* which is the inner-products of *X*. Then we conveniently define an operator 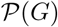 equals to *D*.

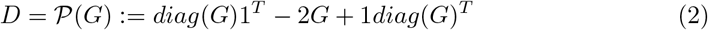

The Gram matrix and the Euclidean distance matrix are linearly related by formula Eq (2). Consider the singular value decomposition of *G*:

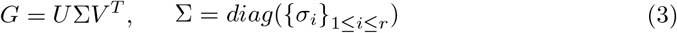

where *U* and *V* are *n × r* matrices with orthogonal columns, and the singular values *σ*_*i*_ are positive. We then have

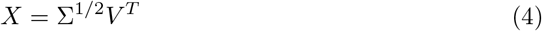

The estimation of the atomic coordinates *X* in the molecule is vital to the NMR techniques for structure determination. We propose a solution to the protein atomic coordinates by recovering the Gram matrix from known entries using optimization framework based on Riemannian measurement.

#### Models

Generally, only some subsets of the distance information can be measured by the NMR experiments. Such a set of data contains important structural information, but it is far insufficient for the complete determination of the structure. The fact that the Gram matrix which is related to distance information is extremely low rank motivates us to apply the matrix completion technique to recover the uncompleted Gram matrix based on very limited observed distance information. This motivates us to apply the MC algorithm in protein structure reconstruction. We first set up a toy model—PDB-deposited-model to check our algorithms’ validity and robustness. After validation, we apply the same algorithms in real NMR proteins deposited in protein data bank(PDB) [24], which we name as NOE-based-model for avoiding confusion. We notice that in protein the distance constraints between the covalent [25] or coplanar atoms [26] only depends on the residue type, and can be always considered as pre-known distances for both models as long as we know the sequence.

In PDB-deposited-model, we extract all the distances between hydrogen atoms within 5*Å* [4] [27] [28] directly from the PDB file, and additionally consider them as pre-known distances. With these distances together with the covalent/coplanar atom distances as known elements in distance matrix, the corresponding PDB-deposited-model is recovered. Obviously, the toy-model contains much more distance information than real NMR experiment and hence is only used to verify the feasibility of the method. In NOE-based-model, we take the NOE distances deduced from NMR experiments instead of the hydrogen distances with 5*Å*. We notice that these NOE distances usually are not the actual but the upper limits. However, in our method we first simply consider them as actual distances in MC stage and then further refine them in the post-processing stage. Different from the PDB-deposited-model, with only the NOE distances and the distance constraints between the covalent or coplanar atoms as known elements in distance matrix, it is still impossible to use the MC algorithm since the distribution of the known distances is too sparse and non-uniform. In MC theory, one can recover most low-rank matrices only when the number of samples *m* obeys *m > Cnrlogn* and they are nearly uniformly distributed [29], where *C* is a positive number (usually *C* is larger than 1, and *n* is the dimension of matrix). In our case, both conditions are violated. To get the condition of MC algorithms satisfied, we add more distance elements from distances estimated by the triangle inequality according to the known elements, for the details see [30].

### Riemannian method

#### formulation

The Gram matrix *G* with its rank *r* can, in principle, be recovered if it is the unique matrix with rank less than or equal to *r* that is consistent with the data [15]. In other words, the low-rank matrix *G* can be solved exactly by the following convex optimisation problem even though the measurable entries have surprisingly small cardinality.

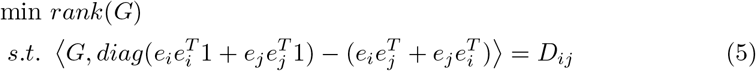

Unfortunately, rank minimization is an NP-hard problem for which the practical solutions take doubly exponential computation time. In order to solve this problem, we follow the work of [22] by rewriting formula Eq (5) as an universal formulation for low-rank MC and adding the boundary constraints:

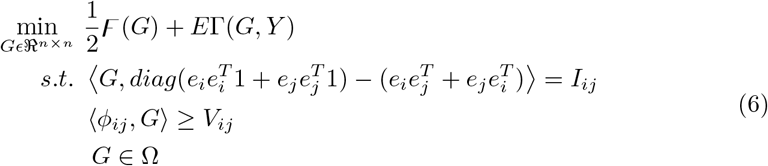

Here, *Y* ∈ ℜ^*n×n*^ is a given matrix, Γ: ℜ^*n×n*^ × ℜ^*n×n*^ → ℜ is a loss function(e.g. *ℓ*_1_ − *loss* function), 𝘍 is a low-rank promoting regularizer, E *>* 0 is the cost parameter and Ω is the structural constraints related to linear subspace. *I*_*ij*_ is an initialized distance matrix, *V*_*ij*_ denotes lower bound restricted by van der Waal’s spheres between pairs of atoms. This type of constraint is designed to prevent structural dislocation from infiltrating each other in non-bonding atoms [31] [32].

Based on the duality theory [33], the solution to problem Eq (6) can be written as *G* = *WW*^*T*^ (*S* + *Q*), where *W* ∈ ℜ^*n×r*^ and *S, Q* ∈ ℜ^*n×n*^.

Then the squared trace norm regularizer is applied to solve the problem Eq (6),

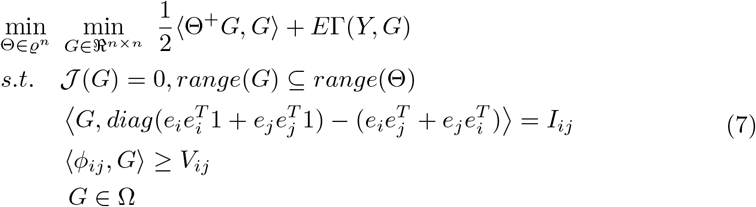

The low-rank constraint on *G* is shifted to 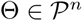 since the ranks of Θ and *G* are equal at optimality [34].

We consider Θ = *WW*^*T*^, and ∥*W*∥_*F*_ = 1. Then the minimized object Eq (7) can be written as follows:

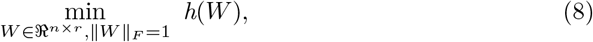

where

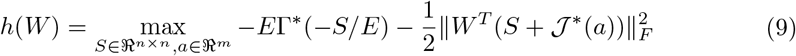

By definition, the gradient of *h*(*W*) above is achieved through the values of variables *S* and *a*. So the recovered matrix *G* is presented as *WW*^*T*^ (*S* + 𝒥*(*a*)). Two effective algorithms are proposed to deal with the problem Eq (9): 1) The Riemannian Cg algorithm: to compute the Riemannian conjugate gradient direction with the step-size by retraction 2) The Riemannian Tr algorithm: to calculate the Riemannian trust-region sub-problem at every iteration [35] [22]. In the end, the local refinement procedure is performed on the Gram matrix, as is usually done in SNL problems [36] [37].

#### Postprocessing

In fact, the Gram matrix after recovered are not completely accurate since the entries sampled from the triangle inequality measurement and NOE experiment have some errors. We perform a postprocessing system to improve accuracy of the rebuilt structures [38].

##### Fixing chiralities

Chirality is an essential factor to discuss the asymmetry in stereochemistry [39] [40]. We perform two types of chirality constraints according to [41]. First, we check the ramachandran angle Φ: if the fraction of positive Φ is larger than 0.5, we simply fix the chirality by adding a negative sign in the x-component of the atom coordinates. Second, every amino acid (except glycine) has two isomeric forms(L- and D-forms). And only L-form is correct enantiomer in life. When the D-form appears at the chiral centers, we fix it by exchanging the coordinates of the group NH_2_ and COOH.

##### BFGS refinement

It is a method for solving the unconstrained nonlinear optimization problems [42]. We employ the functions and parameters in ref [41] to operate the BFGS-based refinement. Using this refinement, we make complements on our MC reconstruction algorithm, where we have treated upper limit as accurate distances in both triangular inequality estimation and NOE distances.

##### EM optimization

It is a process to relax the structure to appropriate bonds and angles. The structure is optimized by minimizing the energy functions of AMBER99SB-ILDN force field [43] in TIP3P water model using simple steepest decent algorithm.

##### Xplor-NIH

We employ Xplor-NIH which is an versatile structure determination and refinement software to improve the resolution of stereostructure [32] [44]. Note that we only use the water refinement part instead of the whole Xplor software here [45].

Finally, we outline the process of reconstructing proteins using the Riemannian method in the form of a flow chart, as shown in Fig 1.

**Fig 1.**
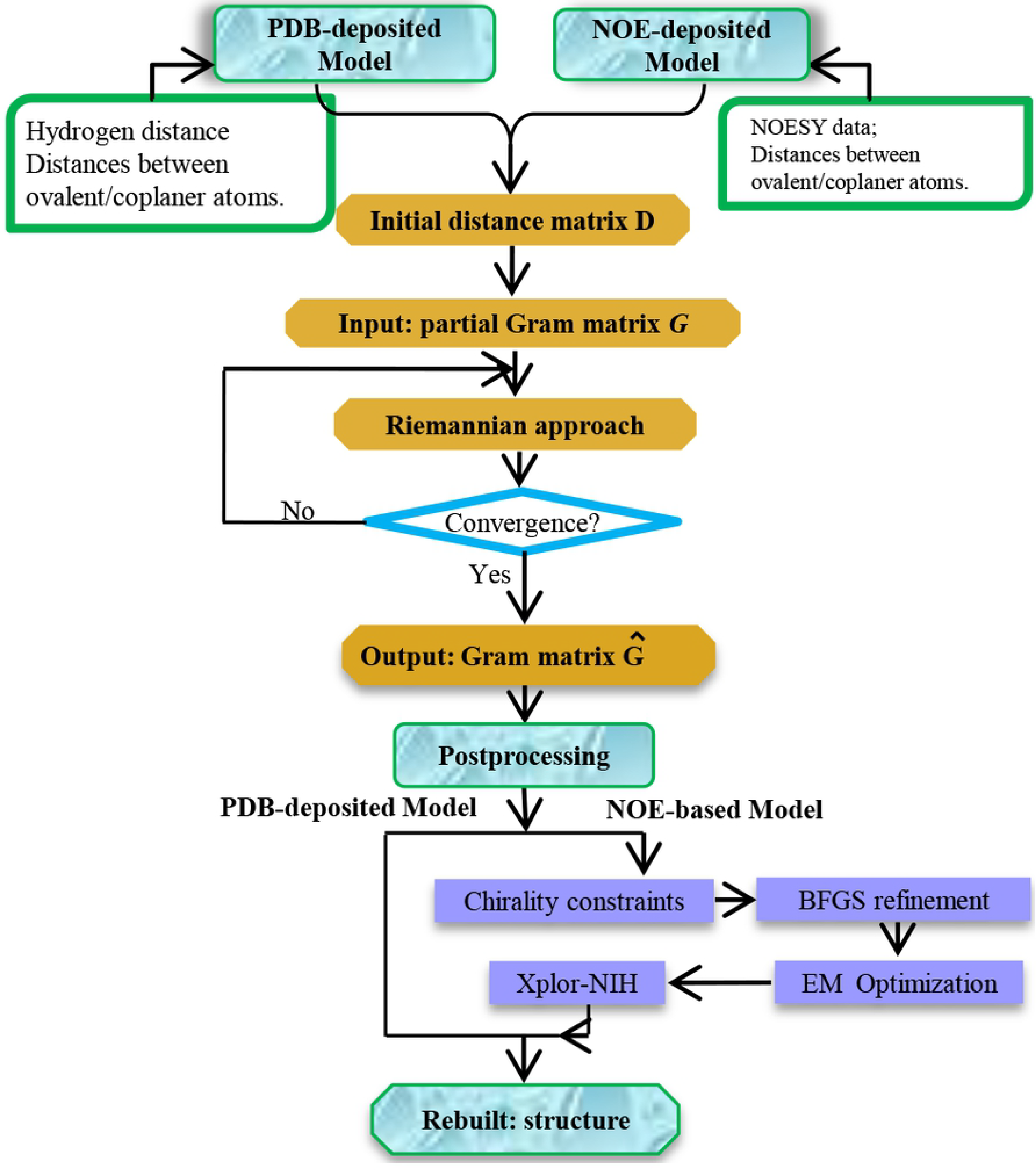
The flow chart of reconstructing the protein structures.

#### Assessment

##### Template Modeling Score (TM-score) and Global Distance Test Score (GDT-TS)

TM-score [46] and GDT-TS [47] were computed to evaluate the structural similarity between protein pairs. According to statistics of structures in the PDB, TM-score below 0.17 corresponds to randomly generated unrelated proteins whereas a score above 0.5 is expected to have the same fold in SCOP/CATH. In general, the higher TM-score and GDT-TS are, the better a rebuilt model is in comparison to reference structure.

##### Protein Structure Validation Software suite

The Protein Structure Validation Software suite (PSVS) [48] is a server to systematically assess and validate the protein structures. PSVS reports knowledge-based quality scores and constraint analyses, such as Z scores, restraint violations, and RMSDs. Z scores contain five geometric validation measures: Verify3D score [49], ProsaII score [50], Molprobity clash-score [51], Procheck Phi-Psi and all dihedral angle G-factor [52]. And higher Z-scores indicate better structures.

##### Ramachandran analyses

Ramachandran Plot is a figure that specifies an enormously allowed conformational region permitted by backbone dihedral angles *ψ*and *ϕ* [53].

##### MolProbity score

MolProbity score (MPscore) combines the clashscore, rotamer, and Ramachandran evaluations into a single score, normalized to be on the same scale as X-ray resolution. The MPscore provides a single value to measure the quality of the prediction structure statistically. Hence, a structure with numerically lower MPscore indicates a more reasonable structure.

## Results and Discussion

50 proteins are picked randomly with different sizes range from 5 to 23 kDa. In the PDB-deposited model, 8 proteins are tested to verify the feasibility of protein structure determination using the Riemannian approach. And in the NOE-based-deposited section, 43 proteins which have both X-ray crystal graphical structure and NMR structure are rebuilt by using Riemannian approach and evaluated with various structure quality assessment metrics. All the coordinate files used are download from PDB database, including reference NMR structures and their X-ray counterparts. The NMR restraints files are extracted in the NMR Restraints Grid of BMRB [54] [55].

### PDB-deposited-model

The detailed recovery results on eight proteins rebuilt by Cg algorithm are shown in Table 1. The performance of Tr algorithm is similar to Cg algorithm, therefore displayed in S1 Table. Table 1 shows that the sampling rate drops sharply with increasing protein atoms *n*. This demonstrates that objects can be perfectly reconstructed from very limited information using Riemannian approach. The backbone RMSDs in well defined (RMSD_bb_wdf) are no more than 1.06*Å* for most of the tested proteins, indicating the rebuilt structures with very high resolution can be achieved using Riemannian approach.

**Table 1.** Reconstruction results of Cg algorithm in PDB-deposited-model

| Entry | Molecule | Seq. | Atoms | Samp. | RMSD_bb_wdf/Cg(Å) |
| --- | --- | --- | --- | --- | --- |
| 5jxv | Immunoglobulin | 56 | 855 | 0.0350 | $0.54 \pm 0.09$ |
| 1g6j | Ubiquitin | 76 | 1228 | 0.0244 | $0.50 \pm 0.07$ |
| 2mx2 | DUB | 81 | 1327 | 0.0226 | $0.29 \pm 0.06$ |
| 2fnb | Fibronectin | 95 | 1416 | 0.0212 | $0.46 \pm 0.07$ |
| 2k49 | UPF | 118 | 1823 | 0.0164 | $0.74 \pm 0.11$ |
| 5h3n | Gelsolin | 133 | 2091 | 0.0143 | $1.02 \pm 0.18$ |
| 5o1t | Protein NRD1 | 179 | 2788 | 0.0108 | $0.32 \pm 0.22$ |
| 6bf2 | Bcl-2-like protein 1 | 212 | 3237 | 0.0093 | $1.06 \pm 0.13$ |

Moreover in order to evaluate the robustness of the method to noisy data, the uncertainty in the distance between hydrogen atoms is analyzed. We assume that the distances between pairs of hydrogen are perturbed by random noises and can be written as follows:

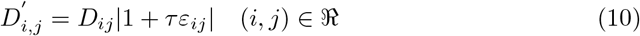

where *D*_*ij*_ is the actual distance between hydrogen atom *i* and *j*. *ε*_*ij*_ ∈ 𝓝(0, 1) is independent standard Normal random variable. The noise is expressed in terms of noise factor *τ*. We implement the low-rank reconstruction on the synthetic data, which simulates the distance data and varies the percentage of the additive noise data.

We take two selected proteins as examples (the other proteins have the similar curve) and present how RMSD_bb_wdf changes over *τ* in S3 Fig. Obviously, it causes increasing RMSD_bb_wdf of Tr or Cg algorithm respectively as the noise factor *τ* grows. The results present that when the RMSD_bb_wdfs of the two tested proteins are equal to 2*Å*, the noise factor values *τ* are 3.09 (2mx2) and 2.16 (2k49) for Tr algorithm, 3.12 (2mx2) and 2.88 (2k49) for Cg algorithm respectively. Meanwhile, the RMSD_bb_wdf of Tr algorithm rises obviously slower than that of Cg algorithm below 4*Å* for the two proteins. This indicates that the Tr algorithm is more robust to the data noise in determining protein structures.

### NOE-based-model

We compare the rebuilt structures using Riemannian approach with NMR structures in terms of metrics on structure similarity, stereochemical quality, restraint violations, and Ramachandran analysis. The NMR structures which are deposited in the PDB are labeled as reference structures.

#### Structure similarity

We take the X-ray structure as the standard structure, and calculate the TM-score/GDT-TS with respect to them for the rebuilt structures and reference structures, respectively. The results are shown in Fig 2, where we can see that among 43 proteins 20 of them reveal higher TM-score/ GDT-TS values (rising an average of 3.48%/3.18%, respectively) for the rebuilt structures with Riemannian approach compared with the reference NMR deposits. In Fig 2 we also see that for most proteins (31/43) Tr algorithm give higher scores than Cg algorithm.

**Fig 2.**
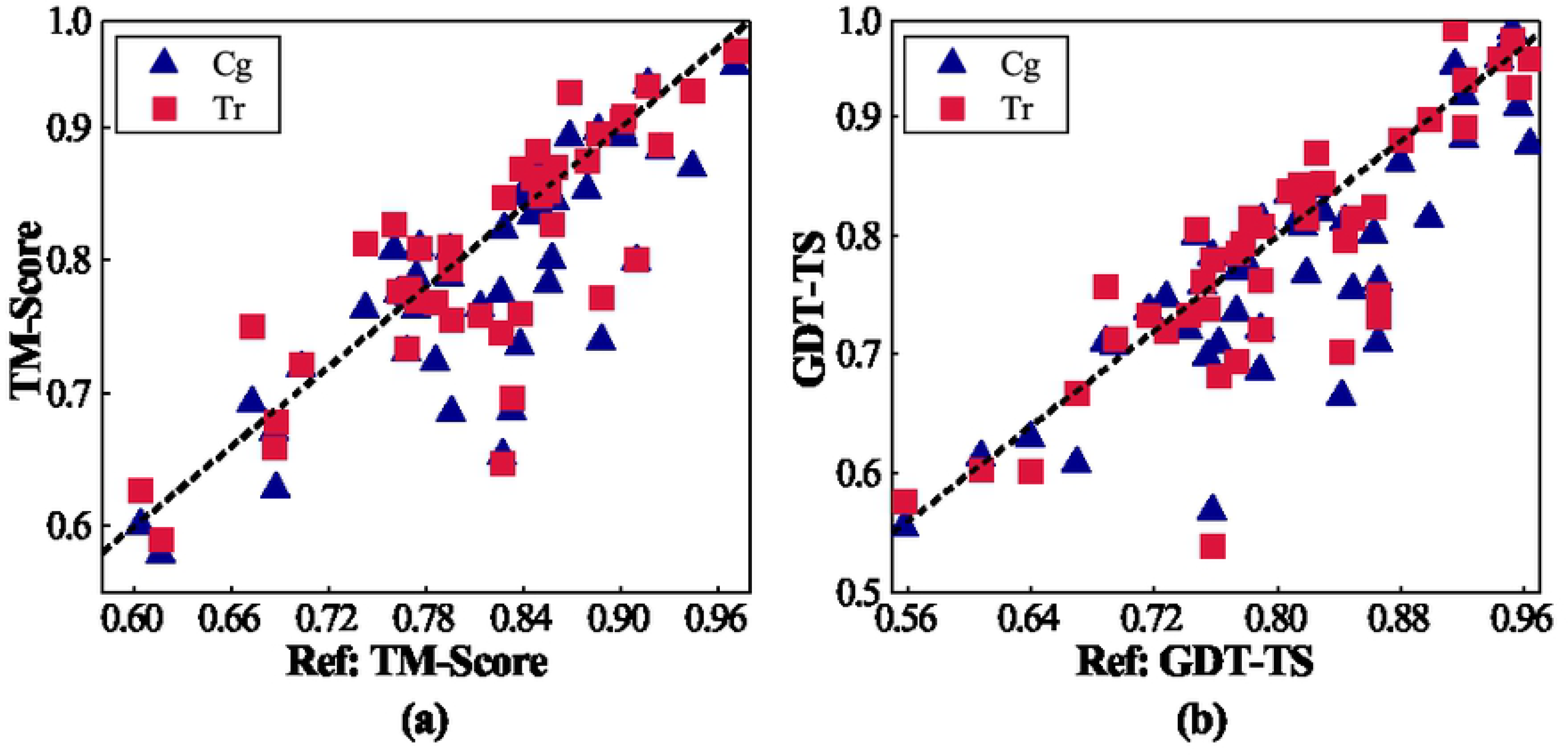
The TM/GDT-TS scores of reference structure and rebuilt structure. (a)/(b) is the TM/GDT-TS score of reference structure and rebuilt structure (with Riemannian approach) using the X-ray counterparts as template. The TM/GDT-TS value of the reference structure is plotted on the X-axis. While the value of Riemannian approach is displayed on the Y-axis. And the oblique lines in (a-d) represent y=x.

The RMSD values of reference structure and rebuilt structure are measured based on corresponding X-ray structure, respectively. We calculate the ratio of the two RMSD values (r_rmsd) to characterize the quality of our reconstructed results. When the ratio is not larger than 1, we say our result is comparable with the PDB reference; otherwise, our reconstruction is worse. The results are shown in Fig 3 and S3 Table. We can see that more than half of the rebuilt structures are comparable or even better in comparison to reference structures. We calculate the average percentage of the pre-known distances for the proteins with r_rmsd<1 and r_rmsd>1, respectively. For the former one, the percentage is about 0.72%±0.24%, while for the latter one it is about 0.56%±0.15%. Hence, we argue that the bad performance for some proteins is due to limited distance measurements. In Fig 4, we select four proteins with r_rmsd<1 and show the superimposition of X-Ray, NMR, and rebuilt structure by Tr algorithm. Clearly, the rebuilt structure appears closer to the X-ray counterparts compared with reference structure. For clarity, we calculate the *Cα* pairwise distances between the reconstructed structure and X-ray counterpart as well as those between the reference structure and X-ray (as shown in Fig 5). we can see that in some region our rebuilt structure are significantly similar to X-ray structure than PDB NMR deposits. These regions may be significant on biological conformation [56].

**Fig 3.**
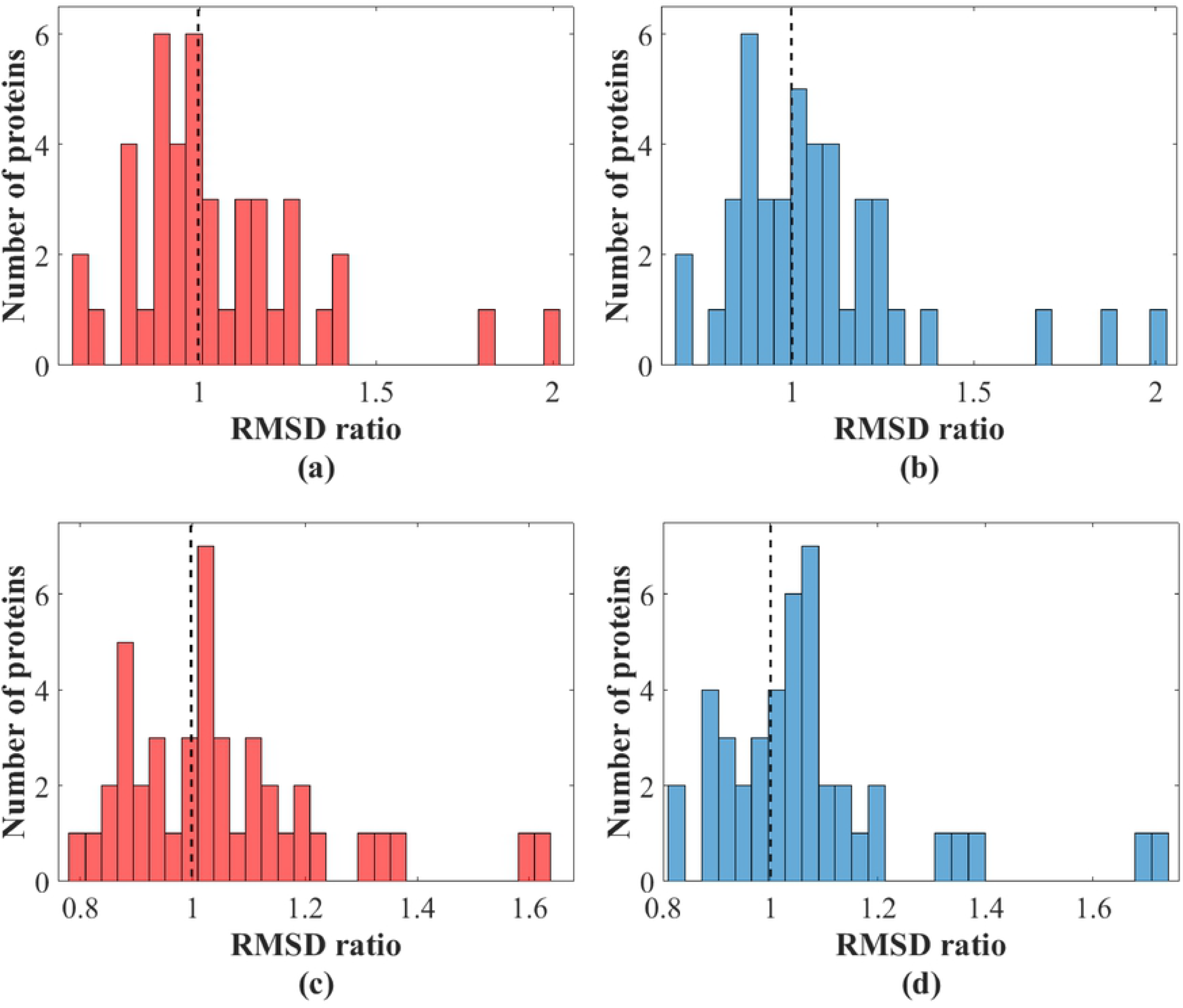
The histogram of backbone and heavy atom RMSD ratio in well-defined region based on X-ray counterpart. The RMSD ratio (r_rmsd) is defined as RMSD_rebuilt/RMSD_reference. The histogram (a) and (c) are rebuilt structures by Tr algorithm, and the Histogram (b) and (d) are rebuilt structures by Cg algorithm.

**Fig 4.**
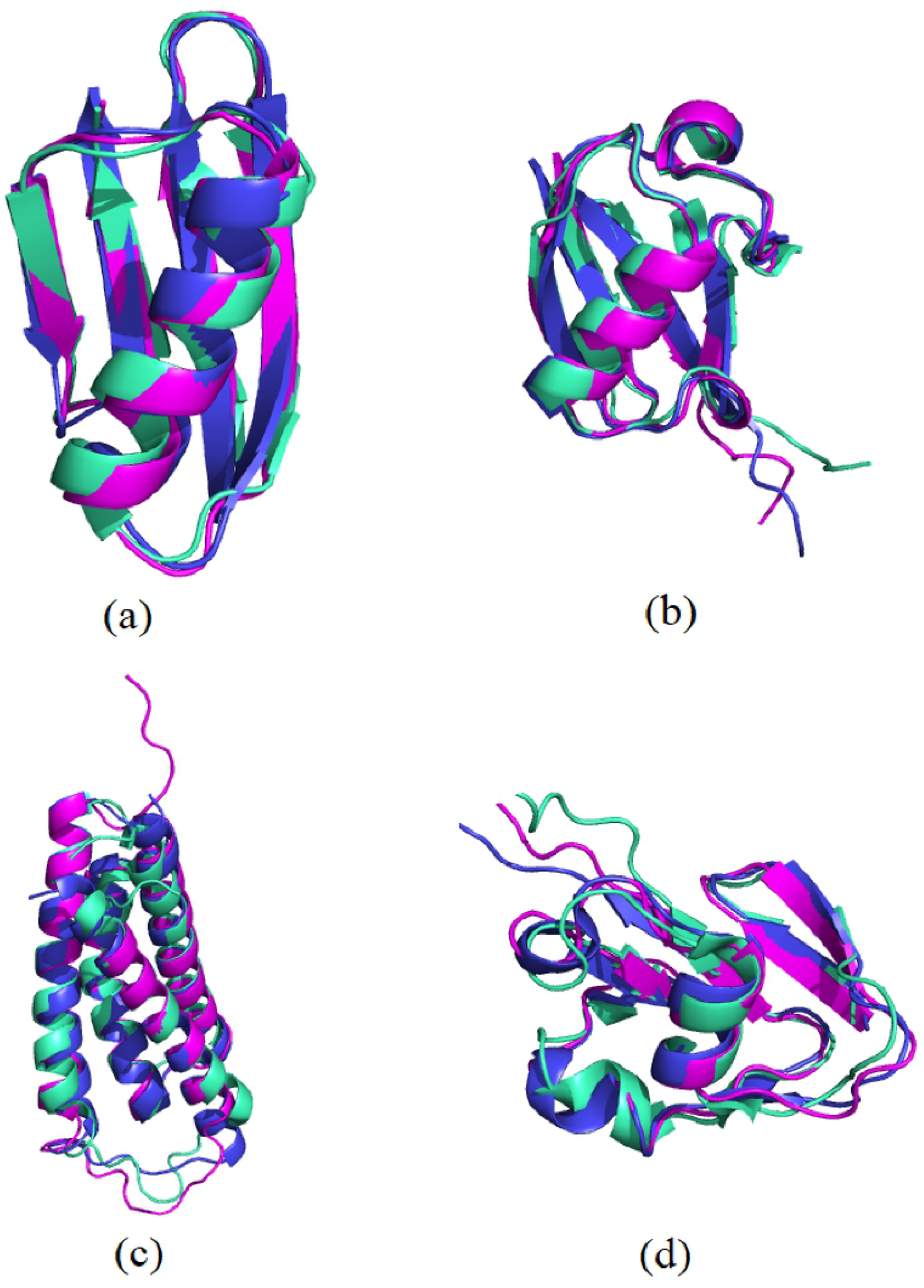
Superimposition of rebuilt structure and reference structures. Superimposition of X-ray crystal structure (blue), reference structure (greencyan), structure rebuilt by Tr algorithm (magenta). The protein entries are (a)-1gb1;(b)-1g6j;(c)-2hfi;(d)-2k5p.

**Fig 5.**
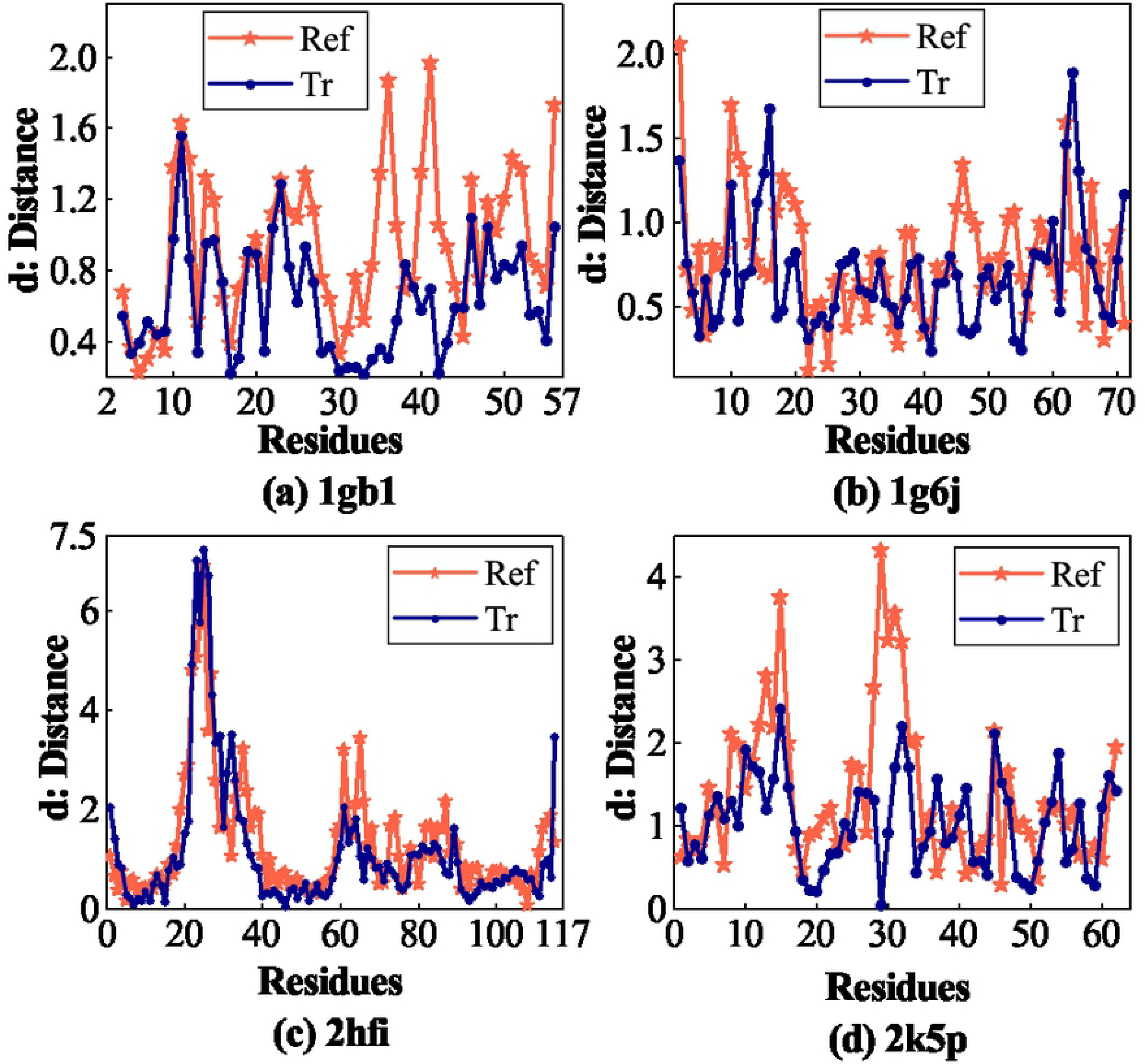
The *Cα* interatomic distance between rebuilt structure/reference structure and corresponding X-ray structure.

#### Stereochemical quality

Five metrics of Z scores are statistically measured by graphical means of boxplot as shown in Fig 6. The average of MolProbity Clash score values using Riemannian approach is almost at parity or slightly lower compared with that of the reference NMR structures, so do ProsaII and Verify3D values. As to Tr algorithm, both Procheck phi-psi and all dihedral angle G-factors are expectedly better than that of the reference structures. This may be due to the help of the dihedral angle and NOE restraints [52].

**Fig 6.**
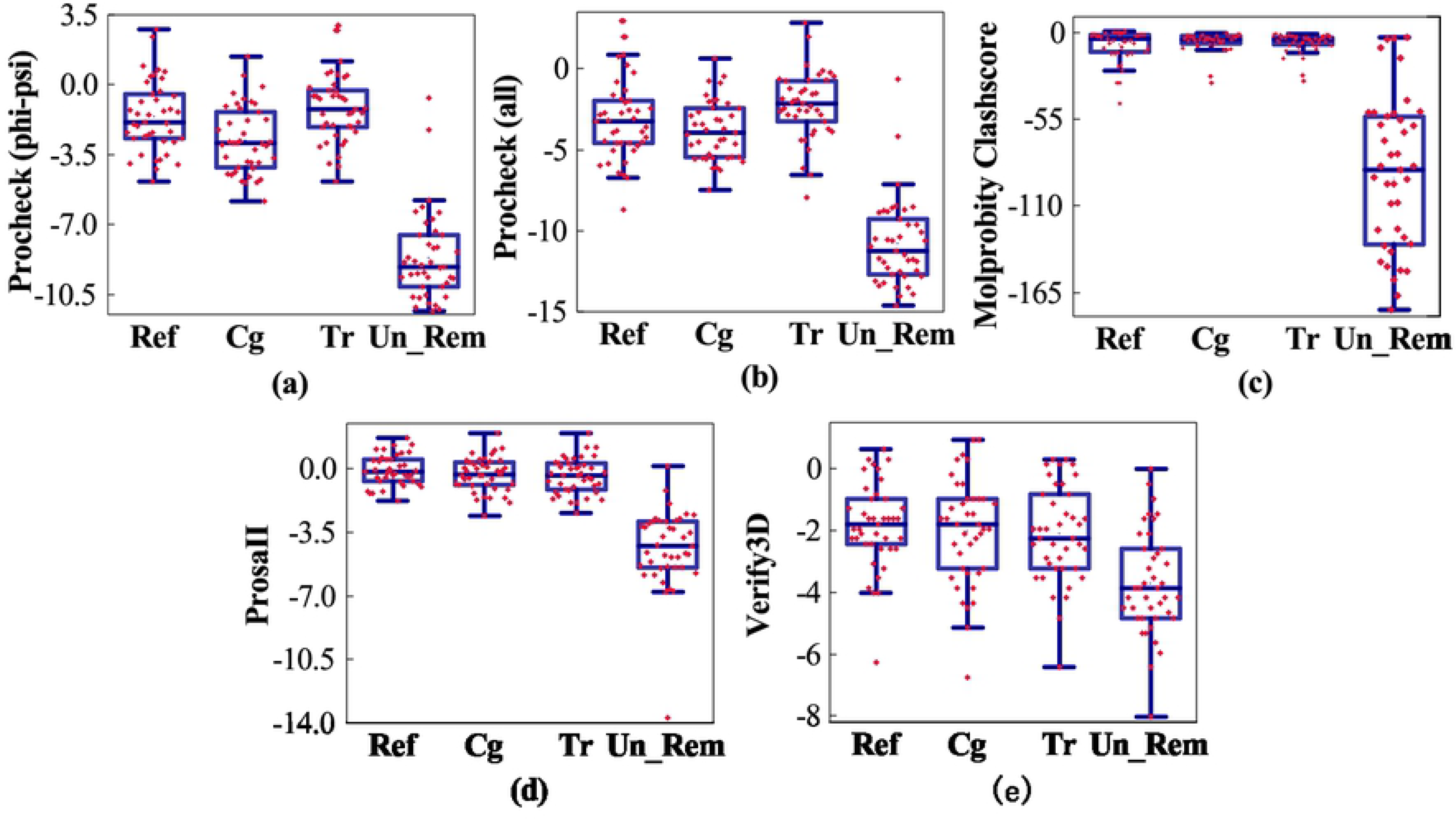
The boxplot of the Z scores for reference structures, structures with and without Riemannian approach, respectively. From (a) to (e), the Z scores orderly are Procheck phi-psi and all dihedral angle, MolProbity Clash score, ProsaII, and Verify3D.

#### Restraint violations

The distance violation and dihedral angle violation are depicted in Fig 7. Distance restraint violation is assessed in terms of ratio (i.e. the number of distance violations divided by the number of distance constraints). Dihedral angle violations per structure are divided into two categories: 1 − 10° and > 10°. The distance violation ratio of rebuilt structure by Riemannian approach is slightly higher than that of the reference structure. However, the maximum distance violation of rebuilt structure using Riemannian approach is lower than that of the reference structure. As to dihedral angle violation, although the performance of the rebuilt structure by Riemannian approach is slightly higher on the 1 − 10° dihedral angle violation, is lower on the > 10° dihedral angle violation compared with the reference structure.

**Fig 7.**
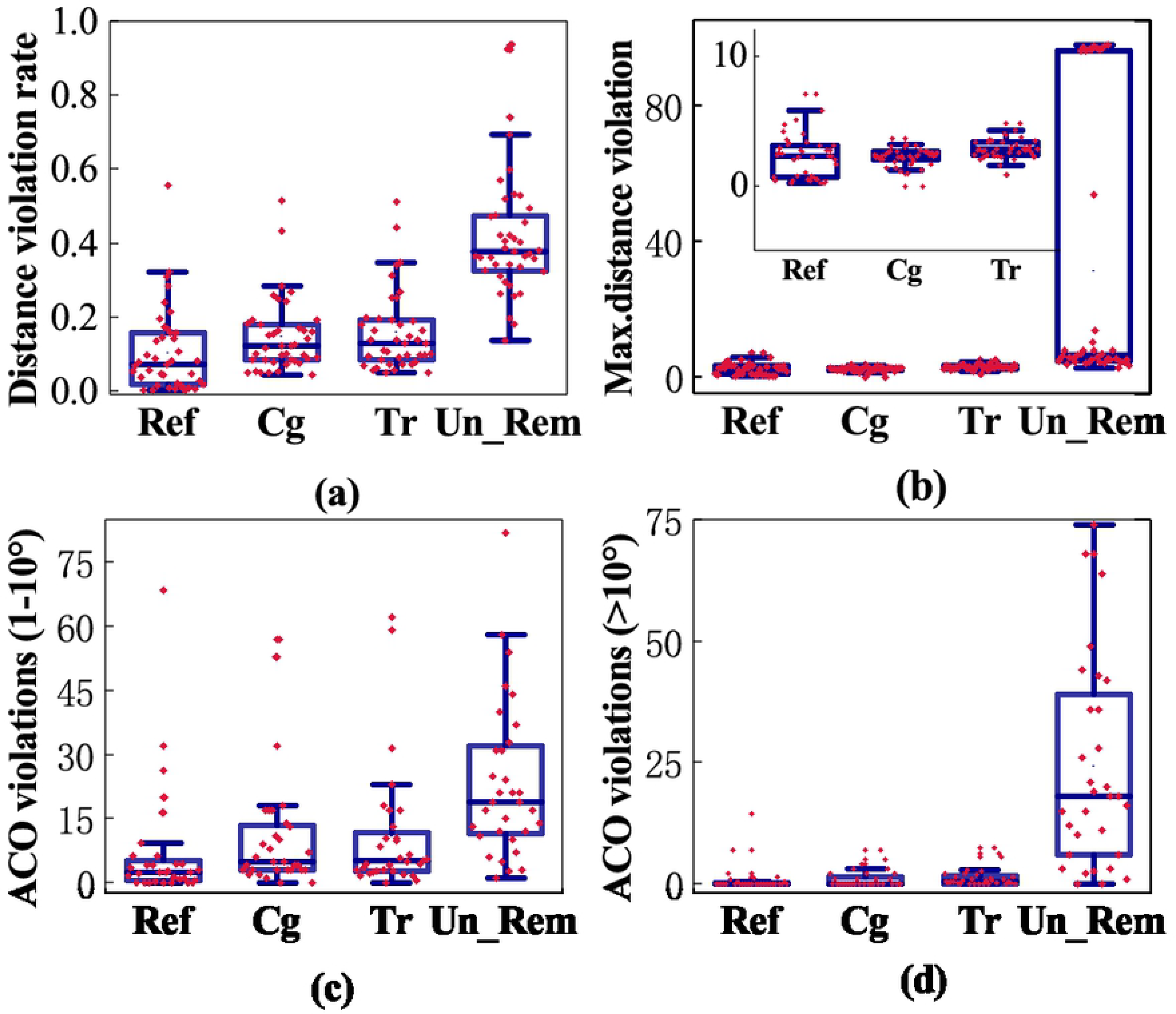
The boxplot of the restraint violations for reference structures, structures with and without Riemannian approach, respectively. The boxplot of (a) is the ratio of the distance restraint violations, (b) is maximum distance violation, (c) is the number of dihedral angle violations between 1° and 10°, and (d) is the number of dihedral angle violations larger than 10°.

#### Ramachandran analysis

Ten reconstructed proteins are picked out to analyze all-atom contacts and geometry [51] for further assessment. As depicted in S1 Fig., the MPscores reduce appreciably to the values less than that of the reference structures with the help of Tr algorithm. And the angle distribution in favored region for tested proteins has a remarkable improvement owing to Tr algorithm-(except for 1r36 and 2hfi).

#### Relation between MC algorithm and post-processing

In the end, we identify the individual contributions of our MC algorithm and post-processing in reconstruction separately. First, we show that with only MC algorithm but without post-processing our reconstructed structure can still have the correct fold(TM score roughly larger than 0.5), but the detailed structure is much worse than the PDB deposits. (as shown in RMSD columns in S2 Table.) Then we illustrate that the MC algorithm is necessary by the following procedure (we labeled this procedure as Un_Rem): we randomly assign the reasonable unknown elements in distance matrix while keeping the known elements intact, and then perform exactly the same post-processing. Then we perform the same assessments on the result of Un_Rem procedure: In S2 Fig., we show the TM-score, where we can see that the Un_Rem method gives very poor results(Even the folds are wrong for most proteins). Besides, we also check the PSVS Z-scores for Un_Rem as shown in Fig 6. Again, the Un_Rem method show very low Z-score indicating the structures from Un_Rem are unreasonable. Hence, we come to the conclusion that the MC algorithm is necessary for get the correct fold of the protein model from the NOE measurements. However, with only MC algorithm, the structure may not be very precise. After proper post-processing, we reduce the errors and improve the quality of reconstructed protein model. As a result, the MC algorithm and post-processing procedure are both important for obtaining high quality protein model from NOE measurement.

Overall, the Tr algorithm in Riemannian approach shows a slight improvement compared with that of Cg algorithm both in quantity and in the magnitude of rise on metrics of structure similarity, stereochemical quality, restraint violations, and Ramachandran analysis. This matches with the analysis of the robustness in PDB-deposited-model: Tr algorithm has better anti-noise performance with the noise factor grows. Not all the reconstructed performance of the tested proteins exceeds the reference structures on all the metrics, but there are still many proteins achieving admirable results. One of the reasons for the worse performance may be that the quantity of the known accurate distance entries is much small. The other may be stemmed from the structural complexity for different proteins. Additionally, the reference NMR structure directly downloaded from the PDB is determined by combining the state-of-the-art methods and perfect refinements. In contrast, the approach we proposed here to determine the protein structure is a general workflow and only used the limited distance geometry information. Even so, there are still many rebuilt structures achieving closer to the X-ray counterparts compared with that of the reference structures. As shown in Table 1 of PDB-deposited-model, the rebuilt structure gains lower RMSD value at so tiny sampling rate. It motivates us to improve the reconstruction results by adding more equality constraints into the distance matrix to solve the problem in NOE-based-model.

## Conclusion

In this paper, the matrix completion technique is presented to the protein structure determination since the Gram matrix, which had linear relation with the protein distance matrix, is extremely low rank. The triangle inequality is introduced to estimate some unknown distance since the known data are too sparse to complete the Gram matrix. The Riemannian approach is proposed and offers two algorithms to recover two models we established. A system of postprocessing is utilized to guarantee the accuracy. The results in the PDB-deposited-model section present high accuracy both for the two algorithms indicating that the Riemannian approach can be use to reconstruct the protein structure. This may be due to the known accurate entries that are enough to rebuild the structures, although these elements are still tiny to the whole matrix. Besides, in NOE-based-model, the rebuilt proteins are assessed from different metrics, such as structure similarity (RMSD and TM/GDT-TS scores), stereochemical quality (Z scores), restraint violations (violations of distance and dihedral angle) and Ramachandran analysis (MPscore and Ramachandran regions). In comparison with the Un_Rem structures, the Riemannian approach achieves a performance that is significantly better on the above metrics. And the Riemannian approach enables for some proteins to more close to their X-ray counterparts than that of reference structures. Our results suggest that the Riemannian approach is a feasible technique to determinate the protein structure and the technique can be expected to obtain high precision results if more distance information is collected.

## Acknowledgments

The authors would like to thank Prof. Choo Hiap Oh for valuable discussions and English modification.

## Supporting information

**S1 Code. The software of protein structure determination based on Riemannian approach.**

**S1 Fig. Ramachandran analysis.** Ramachandran analysis of the reference NMR structures and structures rebuilt by Tr algorithm. And the ten proteins are plotted on the X-axis. (a): the Molprobity score. (b): the angle distribution in favored regions. (c): the outliers of the angle distribution.

**S2 Fig. TM/GDT-TS scores of the rebuilt structure** (a)/(b) is the TM/GDT-TS score of rebuilt structure (with and without Riemannian procedure) using reference structure as template. The TM/GDT-TS value of Tr algorithm is plotted on the X-axis. While the values of Cg algorithm and Un_Rem are displayed on the Y-axis, respectively.

**S3 Fig. Noise analysis.** RMSD errors of the tested proteins with noise parameter *τ* (The value of RMSD_bb_wdf is produced by comparing our rebuilt structures with corresponding Model 1 of reference NMR structure.)

**S1 Table. Reconstruction results of Tr algorithm in PDB-deposited-model.**

**S2 Table. Analysis of rebuilt structure without postprocessing.** The comparison on values of RMSD, TM-score between rebuilt structure by Tr algorithm and reference structure.

**S3 Table. RMSD ratios based on the X-ray structure.** The backbone/heavy atom RMSD ratio of structures between rebuilt structure by Tr algorithm and reference structure.

